## Supplementary Materials for "Weakly supervised classification of aortic valve malformations using unlabeled cardiac MRI sequences"

Fries et al.

### Supplementary Methods

Aortic Valve Localization: we used a threshold-based method to localize the region of the aortic valve. For each MRI sequence $s = \{x_i,\ldots,x_n\}$, we computed a single pixel map of standard deviations across all frames $x_{std} = \sigma(\{x_i,\ldots,x_n\})$. This map was used to compute an Otsu threshold [33] to binarize and label regions with the greatest variation in blood flow. The regions where ranked by weighted area, with the largest region almost always mapping to the aorta. In cases where multiple regions had similar areas (i.e., within 50% of the top ranked candidate), we selected the region closest to the lower left quadrant by doing a secondary sort on centroid y-axis and selected the highest y-value. In a manual confirmation on the development set MRIs (n=106)1, this localization procedure captured the aorta in 95% of cases, with a final cropped size of 32x32 pixels. Each individual frame of this cropped sequence was then binarized, again with an Otsu threshold, to define a noisy segmentation mask for the aorta for the length of the cardiac cycle. All image preprocessing was computed using scikit-image [34].

Cardiac Cycle Alignment: The duration and peak of each cardiac cycle varies across patient and for the MAG and VENC series the aortic valve is only visible during *systole*, i.e., when blood is flowing through the valve. Since valve shape is most visible at peak blood flow, and pixel intensity is a measure of blood flow in MAG images, we identified peak frames by computing the standard deviation of pixel intensities per-frame and selecting the max, $x_{peak} = \max(\sigma(x_1),\ldots,\sigma(x_n))$. Selecting a +/- 7 frame window around this peak captures 99.5% of all pixel variation for the aortic valve. All three sequences are aligned to this peak and window size is treated as a hyperparameter in the models described below.

#### Supplementary Figures

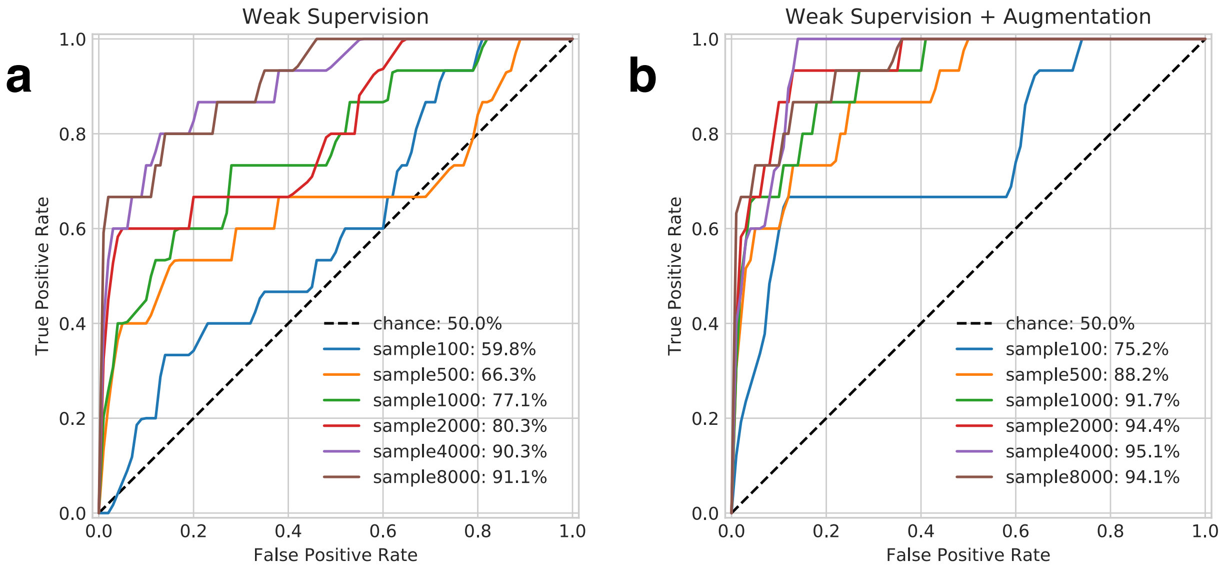


**Supplementary Figure 1. Area under the ROC curve (AUROC) for all scale-up models.** As the CNN-LSTM is trained on more weakly labeled data AUROC generally improves. Plots show models using weak supervision (a) and weak supervision + augmentation (b). In very small training set regimes (e.g., 100 - 1000 instances) using only weakly labeled data, performance degrades after > 0.6 true positive rate.

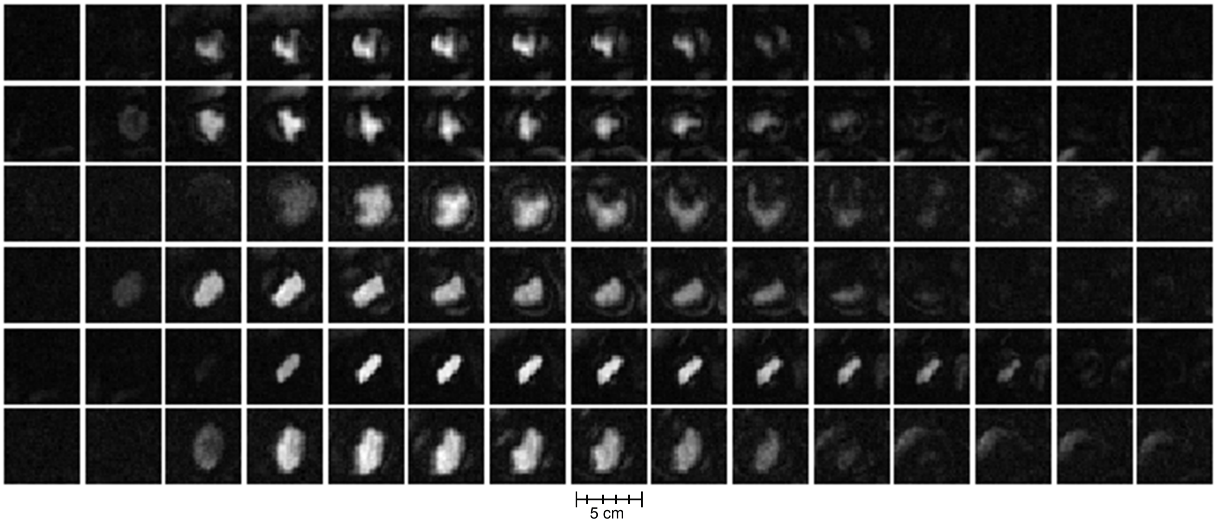


**Supplementary Figure 2. Development set BAV subjects.** All 6 BAV subjects used for labeling function development. For the generative model, 6 contiguous frames performed best at classifying training data using labeling functions, while in the discriminative CNN-LSTM model, 10 frames performed best. This shows how the deep learning model was better able to take advantage of subtle features at the start and end of the cardiac cycle, while labeling functions are restricted to less ambiguous features near the peak frame.

#### Supplementary Tables

##### Supplementary Table 1: Labeling Function Implementation Details

| Name | Heuristic Definition |
|---|---|
| lf_area(x) | -1 if x.area >= 2.13<br>1 if x.area <= 0.99<br>0 otherwise |
| lf_eccentricity(x) | 1 if x.eccentricity >= 0.011<br>-1 if x.eccentricity <= 0.010<br>0 otherwise |
| lf_perimeter(x) | 1 if x.perimeter <= 0.49<br>0 otherwise |
| lf_intensity(x) | 1 if x.intensity >= 2.65<br>-1 if x.intensity <= 2.0<br>0 otherwise |
| lf_ratio(x) | ratio = x.area/x.perimeter**2<br>-1 if ratio >= 4.3<br>1 if ratio <= 4.15<br>0 otherwise |

Python implementations of all labeling functions used in this work. All thresholds were manually set via visual inspection of feature value histograms computed using the development set. By design, we restricted labeling function code to use simple threshold-based heuristics. However labeling functions are black box predictors and can use encode more complex domain heuristics as needed.

**Supplementary Table 2: CNN-LSTM Model Hyperparameter Search Grid**

| **Hyperparameter Name** | **Parameter Grid** |
| --- | --- |
| Learning rate | {1e-3, 1e-4} |
| L2 penalty | {1e-3, 1e-4, 1e-5} |
| Dropout | {0.1, 0.25, 0.5, 0.6, 0.8} |
| Hidden Layer Size | {128, 192, 256} |
| Data Augmentation: Translation | [0.25, 0.25] |
| Data Augmentation: Rotation | [-20, 20] |
| Data Augmentation: Zoom | [0.8, 1.5] |
